# Saliva-based Biomarkers for Predicting Gastric Cancer

**DOI:** 10.1101/2025.05.20.655204

**Authors:** Akanksha Arora, Gajendra Pal Singh Raghava

**Author notes:** **Corresponding author** Prof. Gajendra P. S. Raghava Head and Professor, Department of Computational Biology, Indraprastha Institute of Information Technology, Delhi, Okhla Industrial Estate, Phase III, (Near Govind Puri Metro Station) New Delhi, India – 110020 Office: A-302 (R&D Block), Website: http://webs.iiitd.edu.in/raghava/.

## Abstract

Saliva-based biomarkers offer a non-invasive, convenient, and patient-friendly approach for gastric cancer (GC) diagnosis, eliminating the need for uncomfortable procedures. This study aimed to identify salivary extracellular RNA (exRNA) biomarkers capable of distinguishing GC from normal samples. We used GEO dataset containing exRNA expression profiles of 98 GC and 100 normal samples. We filtered the data and applied the Mann-Whitney U test to identify statistically significant salivary exRNAs (p < 0.05). Initially, a single gene-based classification method identified potential biomarkers including GSDMA, SMR3B, NPTXR, TBCD, KCNC4, and KRT26, achieving accuracies ranging from 61.61% to 64.14%. Then, we applied AI-based methods to identify GC biomarkers. We developed machine learning models on different sets of biomarkers extracted from the original set of statistically significant exRNA using feature selection methods. An optimal primary biomarker set of eight mRNAs (GSDMA, CCDC141, TBCD, STARD13, WRB, ARHGAP23, CDHR3, BX842679.1) was identified, yielding an AUC of 0.905 and MCC of 0.770 with an ensemble stacking classifier. We performed comparative analysis with existing GC biomarker studies which demonstrated that our identified primary biomarker set outperformed previously reported biomarker panels. Out of the eight identified genes, seven have been reported in the literature to be associated with Gastric Cancer. Our results demonstrate the potential of salivary exRNA biomarkers as a non-invasive tool for GC diagnosis.

**Key points:**

- GC is often diagnosed at advanced stages using invasive procedures
- Salivary exRNA has potential for non-invasive GC detection
- A machine learning model was developed to predict GC
- A panel of 8 salivary exRNA biomarkers for GC were identified
- The biomarker panel outperformed existing salivary biomarkers for GC

## Introduction

Gastric cancer (GC) is one of the most common type of cancer accounting for a significant number of cancer-related deaths worldwide [1]. Despite advancements in treatment and management, the prognosis for GC remains poor, primarily due to its late-stage diagnosis [2]. Identifying GC in its early stages is vital for improving survival rates. However, current diagnostic techniques such as upper gastrointestinal (GI) endoscopy with histopathological biopsy are invasive, costly, and impractical for routine screening [3]. Therefore, there is an urgent need for non-invasive and reliable biomarkers that can facilitate early detection and improve patient outcomes. Liquid biopsy has recently emerged as a promising approach for cancer diagnostics, utilizing circulating biomarkers in biofluids such as blood, urine, and saliva [4–6]. Among these, saliva has gained significant attention due to its ease of collection, non-invasiveness, and ability to reflect systemic disease states [7]. Saliva contains a wide range of biomolecules, including extracellular RNA (exRNA) derived which serve as potential biomarkers for various diseases, including gastric cancer [8–10]. Extracellular RNA comprise of mRNA, miRNA, piRNA, miscRNA, circRNA, and other mixed RNA (Figure 1).

**Figure 1:**
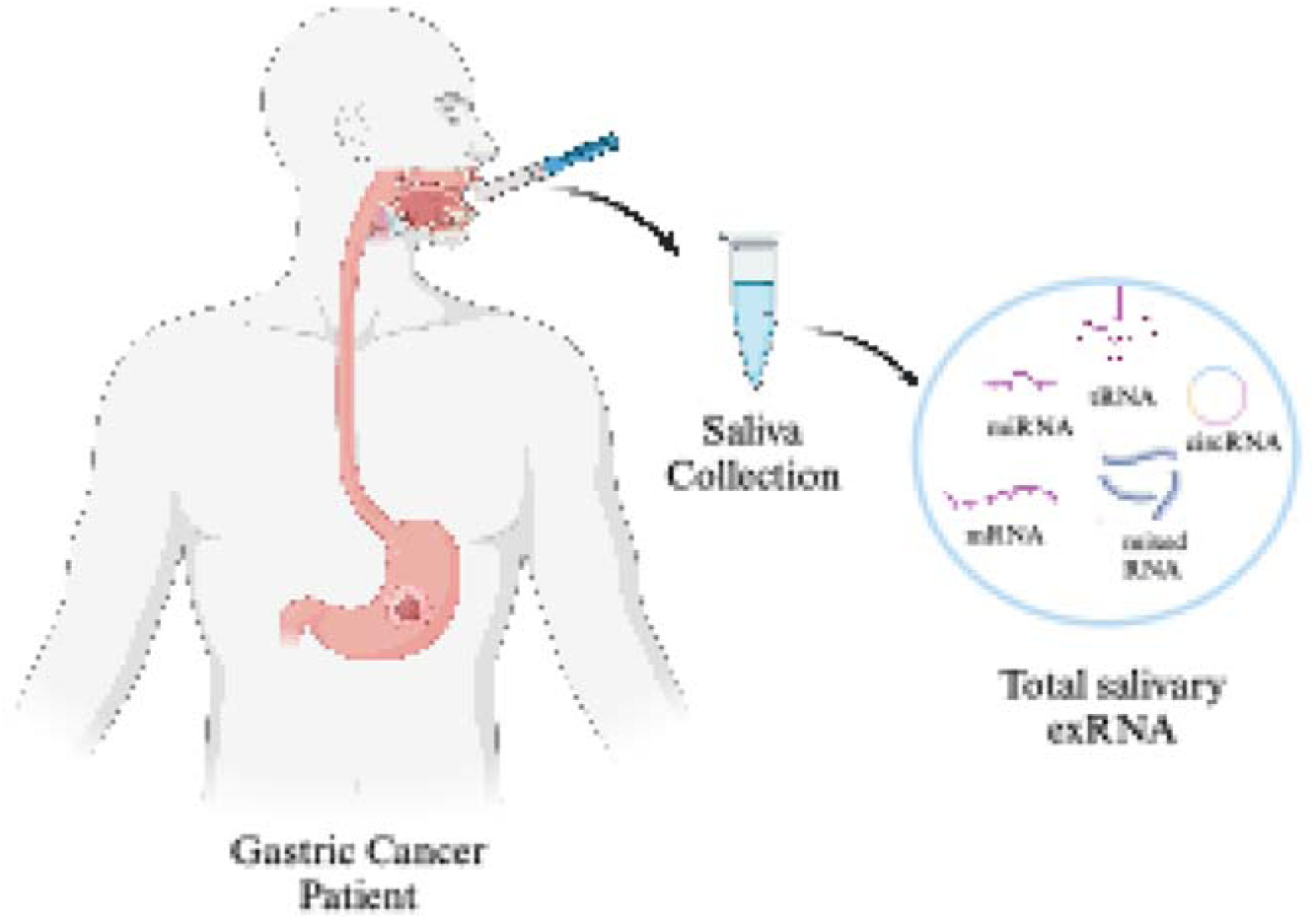
Salivary extracellular RNA profile in GC patients

Various biomarker-based approaches have been explored to improve early detection, utilizing biofluids such as serum, plasma, and saliva. Blood-based biomarkers include carbohydrate antigen 19-9 (CA 19-9), carcinoembryonic antigen (CEA), and pepsinogen levels. However, they exhibit low sensitivity and specificity, particularly in early-stage disease [11]. MicroRNA (miRNA) signatures in plasma have also shown promise, with studies identifying miR-185, miR-20a, miR-210, and miR-92b as potential GC biomarkers, yielding area under the receiver operating characteristic (AUROC) values between 0.65 and 0.75, but require further validation in larger cohorts [12]. A study by Li et al. (2018) investigated the potential of salivary exRNA biomarkers for non-invasive GC detection by profiling saliva samples. The study identified 5 biomarker panel containing SPINK7, SEMA4B, PPL, miR-140- 5p and miR-301a-3p. These were validated using reverse transcription quantitative real-time PCR (RT-qPCR). The final configured biomarker panel achieved an AUC of 0.81, which improved to 0.87 when combined with demographic factors, demonstrating the potential of salivary exRNA for GC screening [9,10].

In this study, we have attempted to classify 100 Normal and 98 GC samples using salivary exRNA biomarkers with high accuracy. We have used single gene-based classification using threshold and machine learning based approaches. We identified 8 mRNA biomarkers that are able to predict Normal vs GC accurately with an AUROC of 0.905 for independent validation sets. To ensure comprehensive biomarker identification, we also explored secondary biomarker sets in addition to the 8 mRNA biomarker panel (primary set). This was done by iteratively removing previously identified markers and reapplying feature selection techniques. This approach allowed us to identify distinct yet relevant biomarker panels with varying levels of correlation, contributing to a more holistic understanding of GC-associated salivary exRNA profiles. The methodology followed in this study is shown in Figure 2. Our findings highlight the potential of salivary exRNA as a promising avenue for non-invasive GC detection, offering an alternative to traditional, invasive diagnostic methods.

**Figure 2:**
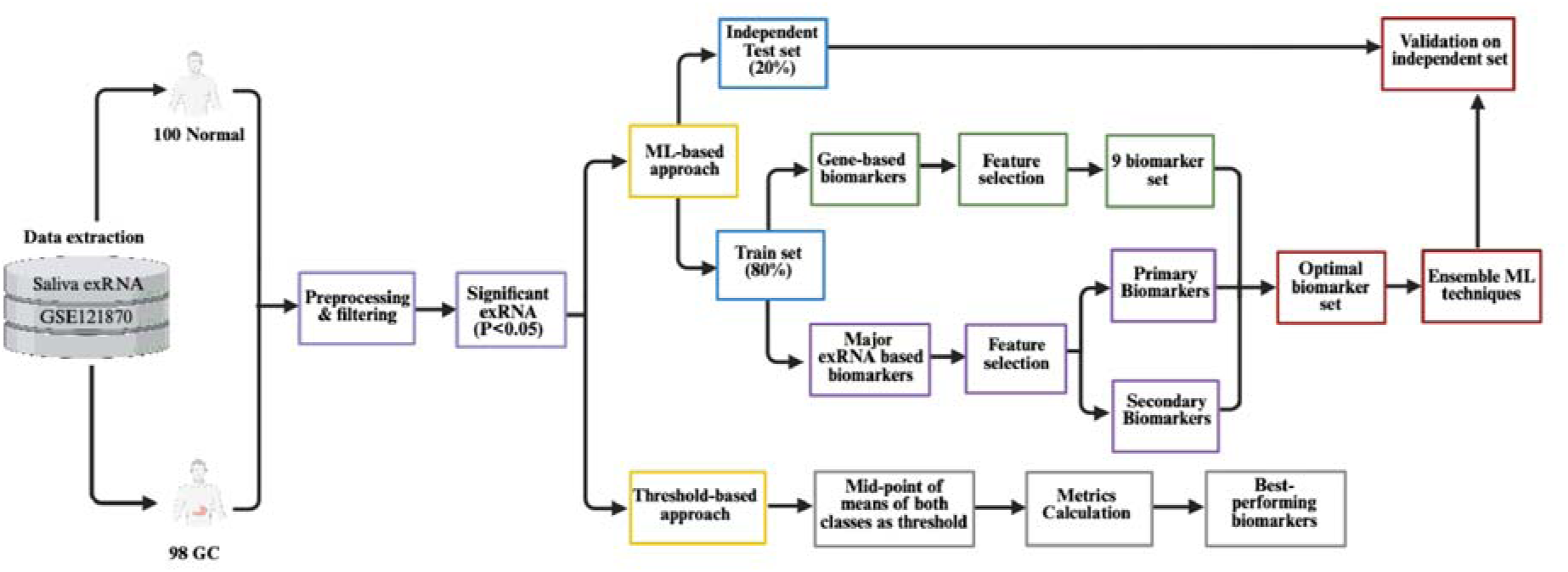
The schematic representation of the methodology

## Methods

### 1. Data Collection

The dataset was collected from Gene Expression Omnibus (GEO) available under GEO accession number GSE121870. It comprises salivary extracellular RNA (exRNA) sequencing data from cell-free saliva samples of 198 individuals, including 100 non-Gastric Cancer controls (Normal) and 98 Gastric Cancer (GC) patients [13]. The dataset had reads in Reads per million (RPM) format for 19704 mRNA, 55104 mixed RNA (lincRNA, snRNA, etc), 1583 miRNA, 2281 piRNA, 26 tRNA, and 245 circRNA.

### 2. Data Preprocessing

The dataset was filtered by removing the exRNA that did not have information for more than 80% of the samples. The RPM values were then transformed using log2 after addition of 1.0 as a constant number to each value. The following equation was used:

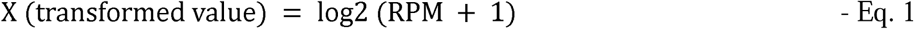

Furthermore, to filter out the statistically significant exRNA, we applied Mann-whitney U test. The Mann-Whitney U test, also known as the Wilcoxon rank-sum test, is a non-parametric test used to compare two independent groups when the data does not follow a normal distribution. After performing the test, we extracted the exRNA that had p-value<0.05. Out of these the most common RNA classes used for biomarkers like mRNA, miRNA, circRNA, piRNA, and tRNA were used for further analysis.

### 3. Threshold-based Approach

In this approach, we attempted to classify the samples into GC and normal using salivary exRNA. For this, we added the means of both the classes and took their mid-point as a threshold to classify GC and normal samples. In this, we computed metrics like True Positives, False Positives, True Negatives, False Negatives, and Accuracy for single biomarkers. We then identified the salivary exRNA having the highest accuracies. These accuracy was calculated as follows:

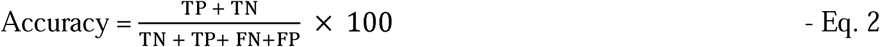

### 4. Machine Learning-based Approach

We applied Machine Learning (ML) models on salivary exRNA considered as features. We used methods like Extra Tree (ET), Random Forest (RF), Logistic Regression (LR), Extreme Gradient Boosting (XGB), k-Nearest Neighbors (KNN), Support Vector Classifier (SVC), AdaBoost Classifier, and Gradient Boost (GB) classifier to classify the samples into GC and Normal [14–21]. In addition to these models, we also applied ensemble machine learning models such as Stacking Classifier (SC) and Voting Classifier (VC) [22–24]. SC combines multiple base models using a meta-model (or meta-classifier) to improve predictive performance by learning how to best combine the outputs of the base models. VC combines predictions from multiple base models by using majority voting for classification to make the final prediction.

#### 4.1. Feature Selection

We used feature selection techniques like Iterative Feature Selection (IFS), Recursive Feature Elimination (RFE), and SVC-L1 to select relevant salivary exRNA that can predict GC and normal samples accurately. RFE progressively eliminates the least relevant features based on model performance, refining the selection to enhance predictive accuracy [25,26]. IFS iteratively adds features, evaluating performance at each step to identify the optimal subset [27]. SVC-L1 leverages a linear Support Vector Classifier with L1 regularization to enforce sparsity, eliminating irrelevant features while preserving key biomarkers for classification [28].

#### 4.2. Gene-based Biomarkers

Firstly, we tried to identify only genes (mRNA) as biomarkers using feature selection techniques, as they were present in the highest amount in the dataset. Gene-based biomarkers are the most commonly studied biomarkers for the diagnosis and prognosis of the diseases.

#### 4.3. Salivary exRNA Biomarkers

Next, we took other classes of RNA into account as well that are commonly used as biomarkers in addition to mRNA, such as miRNA, piRNA, circRNA, and tRNA. These RNA comprised of 362 total significantly different RNAs. We further applied machine learning methods on this set to identify relevant biomarkers.

##### 4.3.1. Primary Biomarkers

We identified a set of primary biomarkers (P) using the best performing feature selection technique - IFS. However, feature selection techniques aim to identify the most relevant features (biomarkers) that contribute to classification, but they often select one representative feature from a group of highly correlated features rather than all of them. This happens because many feature selection methods, such as Recursive Feature Elimination (RFE), Iterative Feature Selection (IFS), and SVC-L1, prioritize reducing redundancy to enhance model interpretability and generalization. In biological datasets, where biomarkers are often co-expressed or functionally related, different feature selection methods might choose slightly different sets of biomarkers depending on their ranking criteria. Thus, there could be multiple relevant biomarkers, but only a subset is selected due to correlations among them.

##### 4.3.2. Secondary Biomarkers

In order to obtain other relevant set of biomarkers, we subtracted the set of identified biomarkers (P) from the total set of salivary exRNA n = 362 (A) leaving us with (A-P) set of salivary exRNA. We then performed the feature selection methods on set (A-P) and performed iterative feature selection again to obtain another set of secondary biomarkers called as S1. We repeated this process again now with the set (A-P-S1), leading to identification of second set of secondary biomarkers S2.

##### 4.3.3. Correlation among biomarkers

To find out if the identified sets of biomarkers – P, S1, and S2, are correlated among each other, we calculated pearson’s correlation for every gene across all three sets. The pearson’s correlation was calculated as follows:

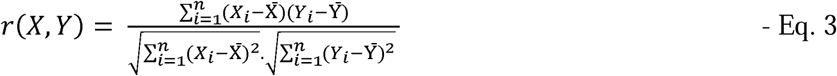

where:

X_i_ and Y_i_ are the expression values of gene X and gene Y in sample i;

X and Y are the mean expression values of gene X and gene Y across all samples

#### 4.4. Cross Validation

To ensure our models remain unbiased and are not prone to overfitting, we implemented a five-fold cross-validation strategy. The dataset was initially divided into training and independent validation sets in an 80:20 ratio. The training data was then subjected to five-fold cross-validation, where it was split into five subsets. In each iteration, four subsets were used for training, while the remaining subset was utilized as a validation set. This process was repeated five times, allowing each fold to serve as the validation set once. The model performance is averaged over the 5 folds to get a robust estimate of generalization, and standard deviation and errors were calculated. The 20% of data set aside during the initial split was used for external validation, providing an independent assessment of the model’s performance. This approach is widely recognized and commonly employed in various studies for rigorous model validation [29,30].

#### 4.5. Performance Metrics

The evaluation of the prediction model is an important step in model development. We have used several threshold-dependent and independent parameters to evaluate our prediction models. The threshold-dependent parameters include sensitivity which calculates how well the model can predict GC (Eq. 4), specificity which calculates correct prediction of Normal subjects (Eq. 5), accuracy which calculated proportion of Normal and GC patients that are correctly predicted (Eq. 6), and Matthew’s correlation coefficient (MCC) evaluated the relationship between observed and predicted values (Eq. 7). The threshold-independent parameter includes AUROC which is a curve plotted between sensitivity and 1-specificity. It reflects the model’s ability to distinguish between different classes. These metrics have been used to evaluate the model in previous studies [31,32].

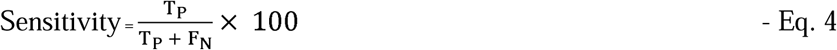

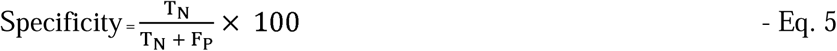

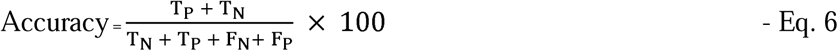

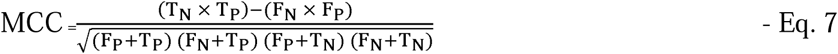

Where T_P_, T_N_, F_P_, and F_N_ stand for true positive, true negative, false positive, and false negative, respectively.

## Results

### 1. Data Preprocessing

The dataset contained 19704 mRNA, 55104 mixed RNA (lincRNA, snRNA, etc), 1583 miRNA, 2281 piRNA, 26 tRNA, and 245 circRNA. After filtering the data, a total of 33212 exRNA which comprised of 11532 mRNA, 516 miRNA, 110 piRNA, 26 tRNA, 10 circRNA, and 21,018 mixed RNA (nonsense mediated decay, lincRNA, snRNA, etc). After applying Mann-Whitney U test, we obtained 1027 exRNA, that were significantly different in both groups with p-value<0.05. The distribution of these 1027 exRNAs is shown in Figure 3 and the total up and downregulated exRNA in each category are given in Table 1. The full distribution of the exRNA categories is shown in Figure 3. Out of these, we used the most common RNA classes of biomarkers like mRNA, miRNA, circRNA, piRNA, and tRNA for further analysis which comprised a total of 362 exRNA.

**Figure 3:**
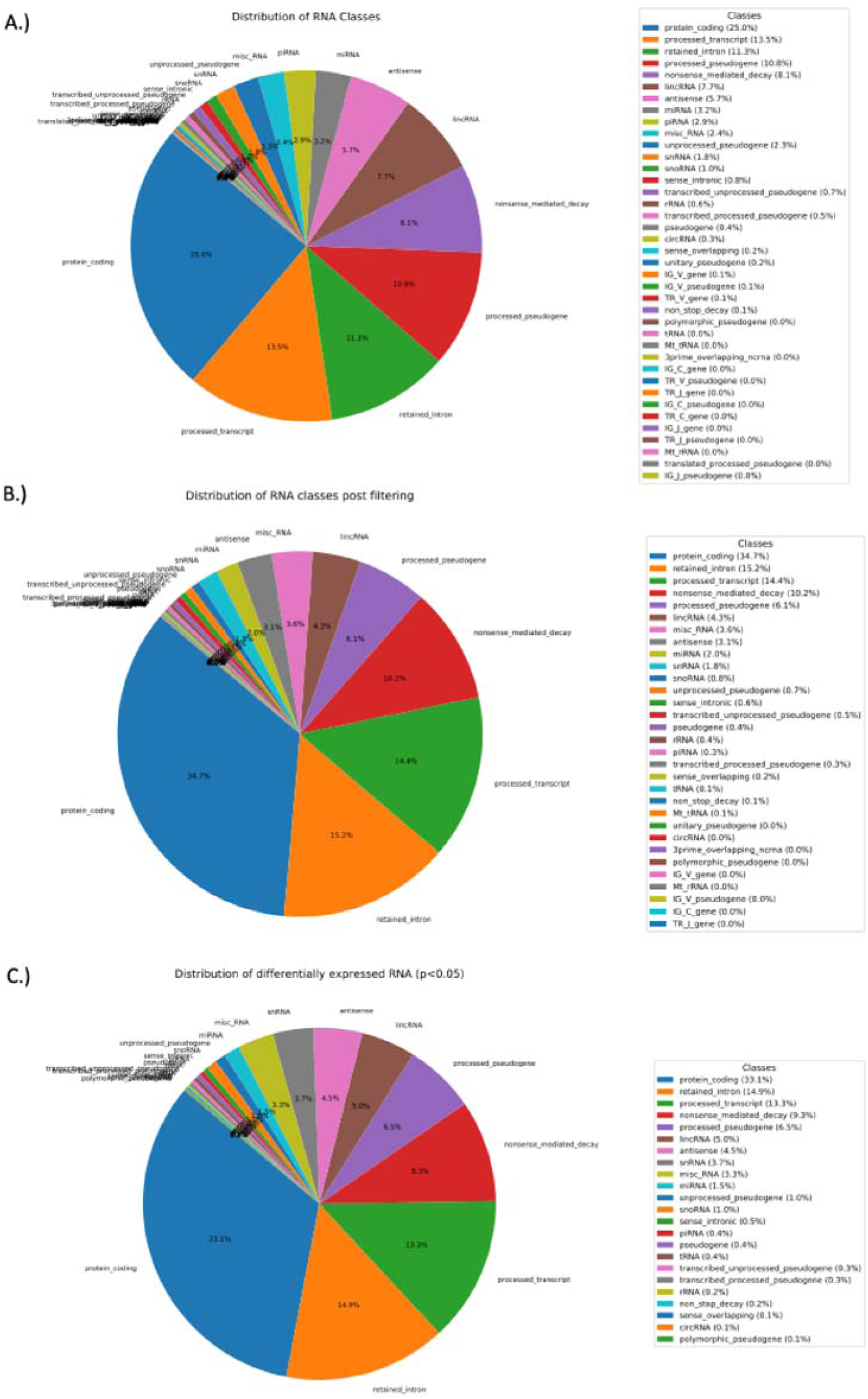
Distribution of RNA classes in A.) the whole dataset, B.) After performing the filtering step: removing of exRNA that had >80% data as zeroes, C.) Significantly different (p-value<0.05) exRNA in GC vs Normal found using Mann-Whitney U test

**Table 1:**
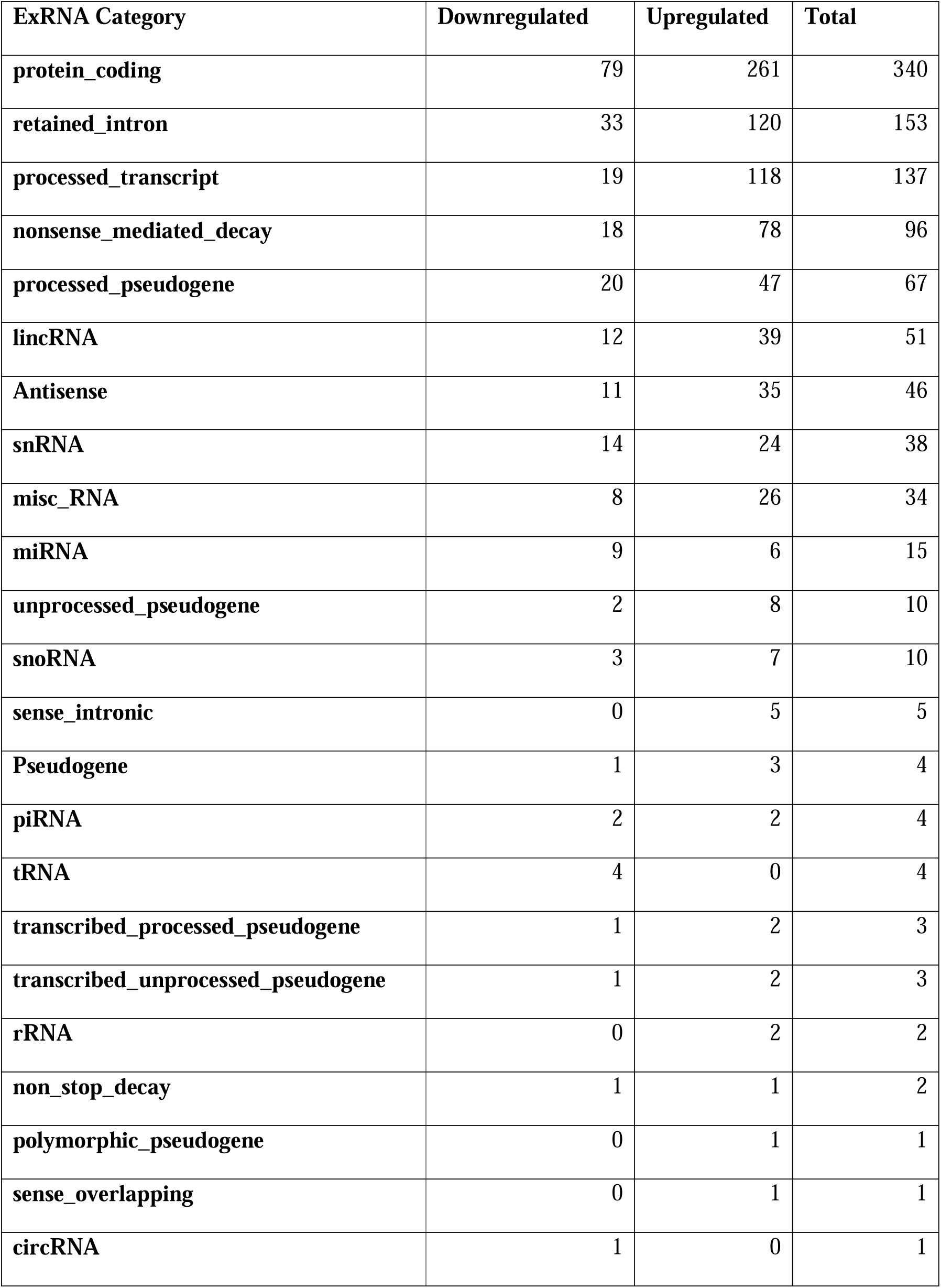
Distribution of upregulated and downregulated exRNAs across different categories.

| ExRNA Category | Downregulated | Upregulated | Total |
| --- | --- | --- | --- |
| protein_coding | 79 | 261 | 340 |
| retained_intron | 33 | 120 | 153 |
| processed_transcript | 19 | 118 | 137 |
| nonsense_mediated_decay | 18 | 78 | 96 |
| processed_pseudogene | 20 | 47 | 67 |
| lincRNA | 12 | 39 | 51 |
| Antisense | 11 | 35 | 46 |
| snRNA | 14 | 24 | 38 |
| misc_RNA | 8 | 26 | 34 |
| miRNA | 9 | 6 | 15 |
| unprocessed_pseudogene | 2 | 8 | 10 |
| snoRNA | 3 | 7 | 10 |
| sense_intronic | 0 | 5 | 5 |
| Pseudogene | 1 | 3 | 4 |
| piRNA | 2 | 2 | 4 |
| tRNA | 4 | 0 | 4 |
| transcribed_processed_pseudogene | 1 | 2 | 3 |
| transcribed_unprocessed_pseudogene | 1 | 2 | 3 |
| rRNA | 0 | 2 | 2 |
| non_stop_decay | 1 | 1 | 2 |
| polymorphic_pseudogene | 0 | 1 | 1 |
| sense_overlapping | 0 | 1 | 1 |
| circRNA | 1 | 0 | 1 |

### 2. Threshold-based Approach

In threshold-based approach, the protein-coding mRNA GSDMA showed the highest accuracy of about 64.14% in classifying the samples into GC and Normal correctly which was downregulated in GC samples, followed by SMR3B which was upregulated and had 62.63% accuracy. The results for top 15 biomarkers are shown in Table 2.

**Table 2:** Results for threshold-based classification of GC and normal samples.

| Biomarkers | TP | FN | TN | FP | Accuracy (%) | Up/Downregulated |
| --- | --- | --- | --- | --- | --- | --- |
| GSDMA | 59 | 39 | 68 | 32 | 64.14 | Downregulated |
| SMR3B | 46 | 52 | 78 | 22 | 62.63 | Upregulated |
| NPTXR | 37 | 61 | 85 | 15 | 61.61 | Upregulated |
| TBCD | 45 | 53 | 77 | 23 | 61.61 | Upregulated |
| KCNC4 | 75 | 23 | 47 | 53 | 61.61 | Downregulated |
| KRT26 | 78 | 20 | 44 | 56 | 61.61 | Downregulated |
| SLC6A11 | 35 | 63 | 86 | 14 | 61.11 | Upregulated |
| KCTD5 | 54 | 44 | 67 | 33 | 61.11 | Upregulated |
| GUCY1A3 | 62 | 36 | 59 | 41 | 61.11 | Downregulated |
| PRKD3 | 42 | 56 | 78 | 22 | 60.61 | Upregulated |
| ZNF250 | 49 | 49 | 71 | 29 | 60.61 | Upregulated |
| SLC30A4 | 52 | 46 | 68 | 32 | 60.61 | Upregulated |
| hsa_piR_019420 | 53 | 45 | 67 | 33 | 60.61 | Downregulated |
| CADPS | 76 | 22 | 44 | 56 | 60.61 | Downregulated |
| PRND | 35 | 63 | 84 | 16 | 60.10 | Upregulated |

### 3. Machine Learning-based Approach

We applied different ML models such as RF, ET, LR, XGB, SVC, GB, KNN, SC, and VC to classify GC vs Normal on different set of features.

#### 3.1. Gene-based Biomarkers

Firstly, we classified the samples into GC and Normal using genes (mRNA), where we selected genes from the initial set of significant exRNA. About 340 significantly different genes were identifies. When we developed ML models on these genes, the maximum AUC achieved was 0.975 on LR model for independent validation set. Next, we identified 9 gene-based biomarkers from a set of 340 genes using feature selection methods. These 9 genes (GSDMA, CCDC141, HSP90B1, SLC30A4, ATP8B3, ARHGAP23, NPTXR, WRB, SMR3B) together achieved the highest AUC of 0.883 on training and 0.893 on independent validation set. The performance of this set is given in Table 3.

**Table 3:** Results for gene-based biomarker sets.

| All significant genes (n=340) |  |  |  |  |  |  |  |  |  |  |  |
| --- | --- | --- | --- | --- | --- | --- | --- | --- | --- | --- | --- |
|  | Training Set |  |  |  |  |  | Independent Validation Set |  |  |  |  |
| Model | Thr | Sens | Spec | Acc (%) | MCC | AUC | Sens | Spec | Acc (%) | MCC | AUC |
| ET | 0.49 | 0.73 | 0.70 | 71.52 | 0.431 | 0.825 | 0.80 | 0.65 | 72.50 | 0.455 | 0.823 |
| RF | 0.51 | 0.77 | 0.79 | 77.85 | 0.557 | 0.846 | 0.80 | 0.70 | 75.00 | 0.503 | 0.845 |
| LR | 0.51 | 0.91 | 0.91 | 91.14 | 0.823 | 0.977 | 0.95 | 0.85 | 90.00 | 0.804 | 0.975 |
| XGB | 0.48 | 0.72 | 0.75 | 73.42 | 0.468 | 0.784 | 0.80 | 0.75 | 77.50 | 0.551 | 0.798 |
| KNN | 0.43 | 0.71 | 0.75 | 72.78 | 0.456 | 0.787 | 0.80 | 0.70 | 75.00 | 0.503 | 0.794 |
| SVC | 0.50 | 0.87 | 0.89 | 87.98 | 0.759 | 0.962 | 0.95 | 0.80 | 87.50 | 0.759 | 0.965 |
| Feature Selection (9 biomarkers) |  |  |  |  |  |  |  |  |  |  |  |
|  |  | Training Set |  |  |  |  | Validation Set |  |  |  |  |
| Model | Thr | Sens | Spec | Acc (%) | MCC | AUC | Sens | Spec | Acc (%) | MCC | AUC |
| ET | 0.49 | 0.83 | 0.80 | 81.65 | 0.633 | 0.861 | 0.75 | 0.80 | 77.50 | 0.551 | 0.860 |
| RF | 0.50 | 0.73 | 0.73 | 72.78 | 0.456 | 0.829 | 0.75 | 0.65 | 70.00 | 0.402 | 0.820 |
| LR | 0.50 | 0.79 | 0.73 | 75.95 | 0.521 | 0.848 | 0.80 | 0.85 | 82.50 | 0.651 | 0.870 |
| XGB | 0.48 | 0.72 | 0.71 | 71.52 | 0.43 | 0.814 | 0.75 | 0.70 | 72.50 | 0.451 | 0.810 |
| KNN | 0.49 | 0.80 | 0.80 | 80.22 | 0.636 | 0.880 | 0.80 | 0.85 | 82.50 | 0.651 | 0.885 |
| SVC | 0.51 | 0.81 | 0.80 | 80.38 | 0.608 | 0.860 | 0.70 | 0.80 | 75.00 | 0.503 | 0.853 |
| Ensemble Classifiers for Feature Selection Set (n=9) |  |  |  |  |  |  |  |  |  |  |  |
|  |  | Training Set |  |  |  |  | Validation Set |  |  |  |  |
| Model | Thr | Sens | Spec | Acc | MCC | AUC | Sens | Spec | Acc | MCC | AUC |

**Table 3:** Results for gene-based biomarker sets
|  |  |  |  | (%) |  |  |  |  | (%) |  |  |
| --- | --- | --- | --- | --- | --- | --- | --- | --- | --- | --- | --- |
| **SC (Base: SVC, ET, LR, KNN, GNB; Meta: KNN)** | 0.48 | 0.81 | 0.80 | 80.38 | 0.608 | 0.883 | 0.90 | 0.80 | 85.00 | 0.704 | 0.893 |
| **VC (SVC, ET, KNN, GNB)** | 0.53 | 0.81 | 0.81 | 81.01 | 0.620 | 0.871 | 0.85 | 0.80 | 82.50 | 0.651 | 0.870 |

#### 3.2. Salivary exRNA Biomarkers

Next, we identified biomarkers from other salivary exRNA categories as well that are commonly used in addition to mRNA, such as miRNA, piRNA, circRNA, and t-RNA. These categories together accounted for 362 exRNA that were significantly different between the two classes. To classify GC and Normal samples, we firstly used the whole set of features and achieved the highest AUROC of 0.97 for training set and 0.965 for the independent validation set for LR model (Table 4). Since, this is set contains very high number of biomarkers and we wanted to identify top biomarkers that can classify GC vs Normal accurately, we applied feature selection to this set.

**Table 4:**
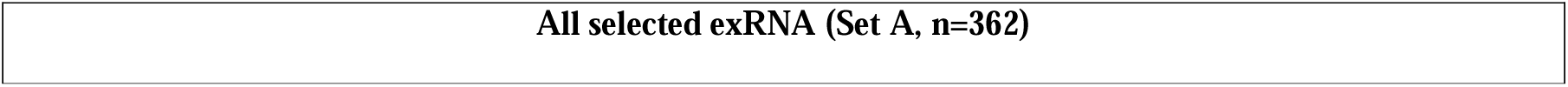
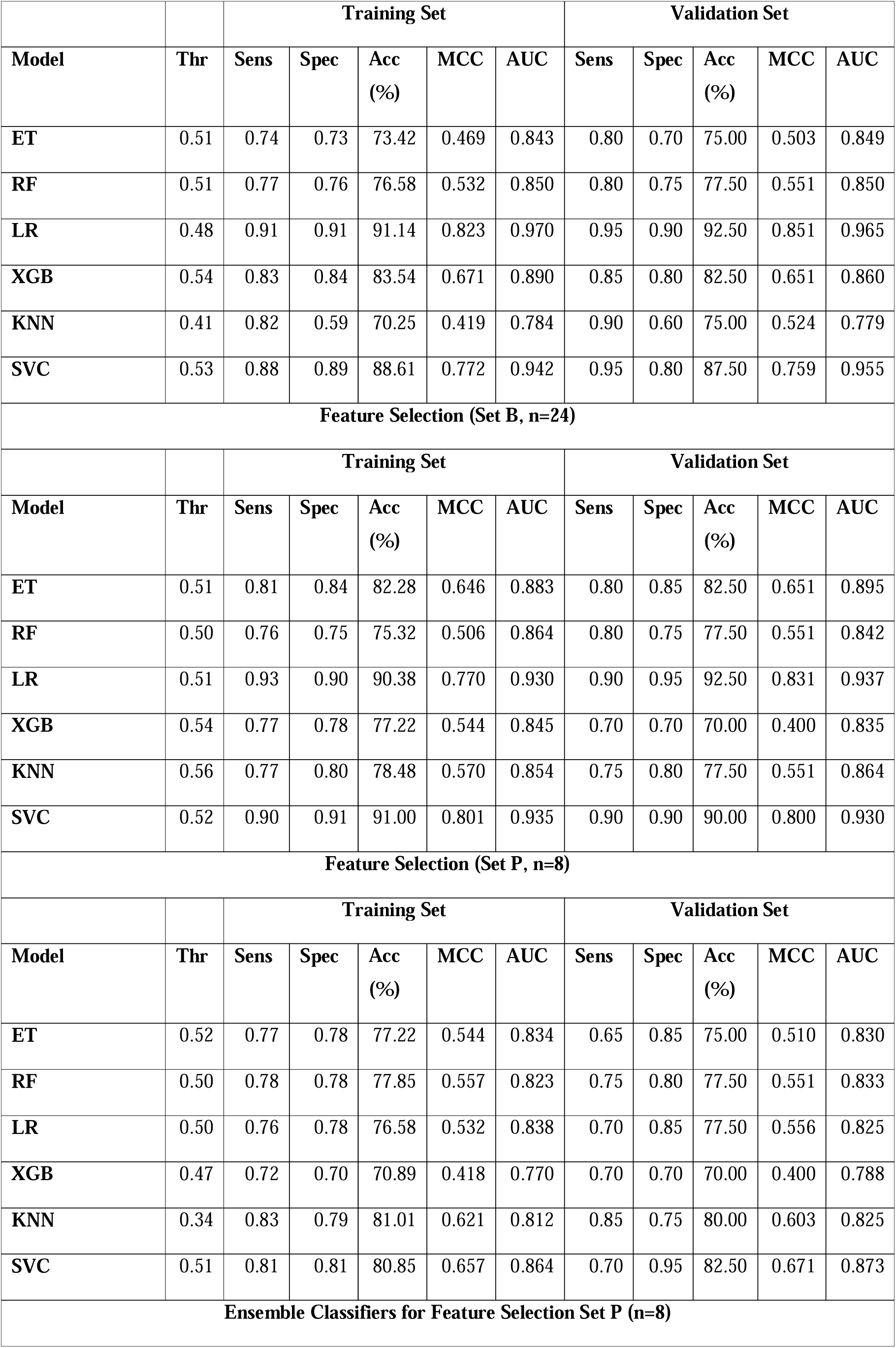
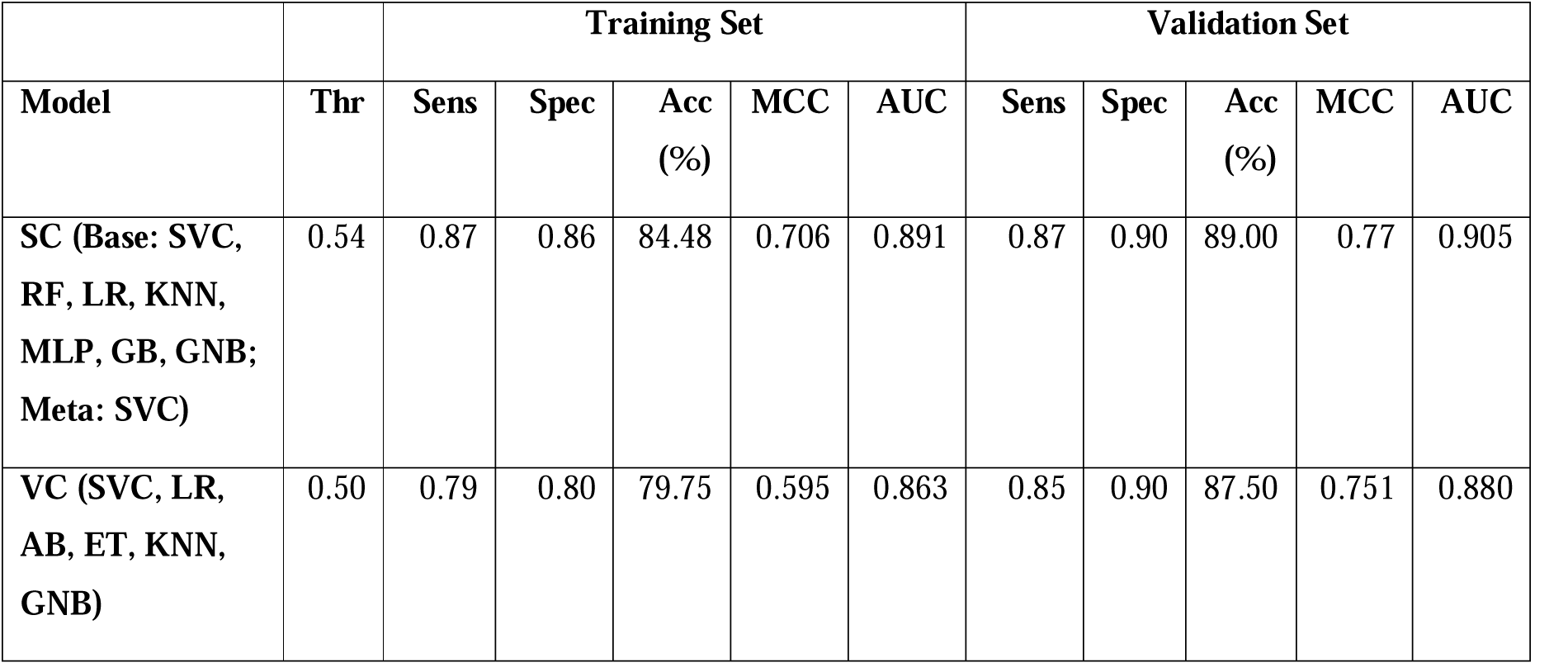
Results on different set of features after feature selection – a) n = 362, b) n = 24, and c) n = 8; here, Thr: threshold, Sens: sensitivity, Spec: specificity, AUC: area under the curve, MCC: matthew’s correlation coefficient.

##### 3.2.1. Primary biomarkers

It was observed that the performance kept decreasing as we reduced the number of biomarkers. However, we were able to achieve comparable performance at 24 features (Set B) using IFS which contained 23 mRNA (genes) and 1 tRNA, leading to the highest AUROC of 0.937 on the independent validation set. We further applied feature selection techniques to identify the top performing biomarkers. Using feature selection techniques, we identified top 8 biomarkers (GSDMA, CCDC141, TBCD, STARD13, WRB, ARHGAP23, CDHR3, BX842679.1) which were able to achieve the highest AUC of 0.873 on independent validation set for SVC model, which further increased to 0.905 when an ensemble method called stacking classifier was applied. We named this best-performing set as primary biomarkers set (P). This primary set of biomarkers contained genes as well, however, it exceeded the performance of the previous set. The detailed results for primary biomarkers are shown in Table 4 and AUROC plots are shown in Figure 4.

**Figure 4:**
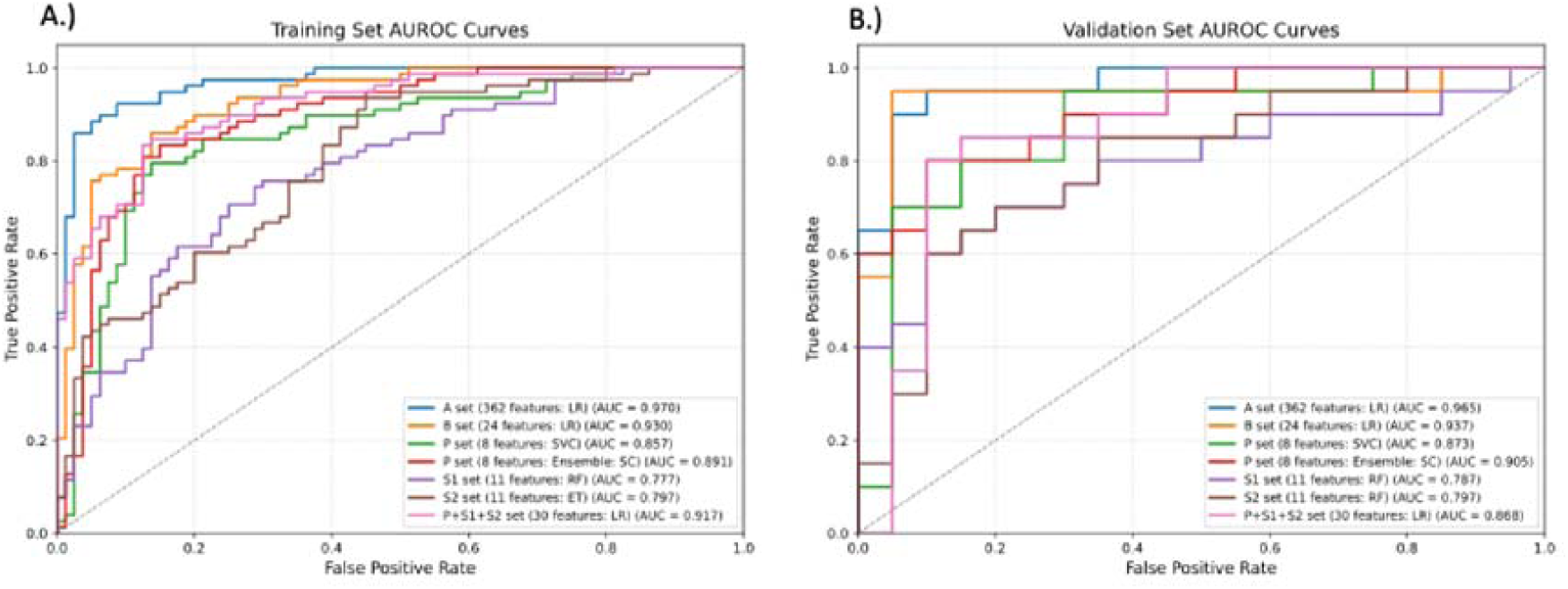
AUROC plots for Training set and Independent Validation set. Here, Set A: All significant biomarkers (n=362), Set B: Biomarkers selected through feature selection (n=24), Set P: Primary set of biomarkers after feature selection (n = 8), Set S1 and S2: First (n=11) and second (n=11) sets of secondary biomarkers

##### 3.2.2. Secondary biomarkers

As mentioned in section 4.1.1, it is seen that feature selection techniques tend to select one representative feature out of the multiple correlated ones, leaving out some biomarkers that can also be used for the classification. For this, we attempted to identify other set of biomarkers as well called as secondary biomarkers by removing the primary set of biomarkers from the total set (n=362). We identify sets S1 (n=11) (KRT26, GUCY1A3, KCTD5, CADPS, ATR, ALX1, TMEM98, HIST1H3A, KPTN, hsa_circ_000353, ZNF688) and S2 (n=11) (KIAA2022, PABPC3, MRPL51, CALB2, MDH1B, ZNF277, PRKD3, EXOSC6, LPPR5, ATP8B3, ZMAT5) as described in section 4.1.2. The RF model developed on secondary biomarker set S1 was able to achieve the highest AUROC of 0.788 on an independent validation set, and the ensemble model (voting classifier) developed on secondary biomarker set S2 achieved the highest AUROC of 0.798 for independent validation set. We added these secondary biomarkers to primary biomarker set to see if the performance is increased after adding them. However, the highest AUC for the combination of these sets (P+S1+S2) for independent validation set was 0.868 using LR classifier. The complete results are shown in Table 5 and AUROC plots are shown in Figure 4.

**Table 5:** Results for Secondary feature sets (S1, S2), and Secondary feature sets S1 and S2 combined with Primary feature set (P)

| Secondary Biomarkers (Set S1, n=11) |  |  |  |  |  |  |  |
| --- | --- | --- | --- | --- | --- | --- | --- |
|  |  | Training Set |  |  |  |  | Validation Set |

| Model | Thr | Sens | Spec | Acc | MCC | AUC | Sens | Spec | Acc | MCC | AUC |
| --- | --- | --- | --- | --- | --- | --- | --- | --- | --- | --- | --- |
| ET | 0.52 | 0.68 | 0.71 | 69.62 | 0.392 | 0.771 | 0.60 | 0.80 | 70.00 | 0.408 | 0.768 |
| RF | 0.53 | 0.70 | 0.71 | 70.35 | 0.397 | 0.777 | 0.75 | 0.70 | 72.50 | 0.451 | 0.788 |
| LR | 0.51 | 0.69 | 0.69 | 68.99 | 0.380 | 0.782 | 0.70 | 0.66 | 67.50 | 0.351 | 0.752 |
| XGB | 0.53 | 0.65 | 0.66 | 65.82 | 0.316 | 0.746 | 0.75 | 0.55 | 65.00 | 0.306 | 0.742 |
| KNN | 0.43 | 0.79 | 0.63 | 70.89 | 0.426 | 0.742 | 0.65 | 0.60 | 62.50 | 0.250 | 0.743 |
| SVC | 0.48 | 0.69 | 0.69 | 68.99 | 0.380 | 0.759 | 0.70 | 0.65 | 67.50 | 0.350 | 0.750 |
| **Secondary Biomarkers (Set S2, n=11)** |  |  |  |  |  |  |  |  |  |  |  |
|  |  | **Training Set** |  |  |  |  | **Validation Set** |  |  |  |  |
| Model | Thr | Sens | Spec | Acc | MCC | AUC | Sens | Spec | Acc | MCC | AUC |
| ET | 0.5 | 0.68 | 0.68 | 67.72 | 0.354 | 0.797 | 0.75 | 0.70 | 72.50 | 0.451 | 0.798 |
| RF | 0.49 | 0.68 | 0.68 | 67.72 | 0.354 | 0.785 | 0.70 | 0.65 | 67.50 | 0.350 | 0.788 |
| LR | 0.48 | 0.69 | 0.69 | 68.99 | 0.380 | 0.759 | 0.65 | 0.65 | 65.00 | 0.300 | 0.735 |
| XGB | 0.47 | 0.69 | 0.69 | 68.99 | 0.380 | 0.748 | 0.80 | 0.70 | 75.00 | 0.503 | 0.744 |
| KNN | 0.51 | 0.68 | 0.73 | 70.25 | 0.405 | 0.741 | 0.75 | 0.55 | 65.00 | 0.306 | 0.734 |
| SVC | 0.49 | 0.72 | 0.70 | 70.89 | 0.418 | 0.781 | 0.75 | 0.60 | 67.50 | 0.354 | 0.771 |
| **Primary + Secondary Biomarkers ( Sets P+S1+S2, n=30)** |  |  |  |  |  |  |  |  |  |  |  |
|  |  | **Training Set** |  |  |  |  | **Validation Set** |  |  |  |  |
| Model | Thr | Sens | Spec | Acc | MCC | AUC | Sens | Spec | Acc | MCC | AUC |
| ET | 0.5 | 0.81 | 0.83 | 81.65 | 0.633 | 0.893 | 0.85 | 0.75 | 80.00 | 0.603 | 0.857 |
| RF | 0.53 | 0.76 | 0.76 | 75.95 | 0.519 | 0.850 | 0.75 | 0.80 | 77.50 | 0.551 | 0.863 |
| LR | 0.51 | 0.85 | 0.85 | 84.81 | 0.696 | 0.917 | 0.85 | 0.70 | 77.50 | 0.556 | 0.868 |
| XGB | 0.48 | 0.79 | 0.80 | 79.75 | 0.595 | 0.851 | 0.70 | 0.80 | 75.00 | 0.503 | 0.830 |
| KNN | 0.61 | 0.71 | 0.81 | 75.95 | 0.521 | 0.821 | 0.60 | 0.95 | 77.50 | 0.587 | 0.826 |
| SVC | 0.48 | 0.78 | 0.79 | 78.48 | 0.570 | 0.869 | 0.75 | 0.80 | 77.50 | 0.551 | 0.859 |

##### 3.2.3. Correlation among biomarkers

We calculated Pearson’s correlations among biomarkers in set P, S1, and S2. The top 3 highest correlated biomarkers for each set are given in Table 6. The highest correlated biomarkers for set P and S1 were GDSMA with GUCY1A3 with a correlation of 0.528, whereas for sets P and S2 – biomarkers ARHGAP23 and LPPR5 were most correlated with correlation of 0.411. For sets S1 and S2, KRT26 and ZNF277 were most correlated with correlation of 0.365.

**Table 6:** Top 3 correlated biomarkers among sets P (Primary Set), S1 (Secondary Set 1), and S2 (Secondary Set 2)

| Top 3 correlated biomarkers in sets P and S1 |  |  |
| --- | --- | --- |
| Biomarker (P) | Biomarker (S1) | Correlation |
| GSDMA | GUCY1A3 | 0.529 |
| GSDMA | KRT26 | 0.404 |
| GSDMA | hsa_circ_000353 | 0.327 |
| Top 3 correlated biomarkers in sets P and S2 |  |  |
| Biomarker (P) | Biomarker (S2) | Correlation |
| ARHGAP23 | LPPR5 | 0.411 |
| TBCD | EXOSC6 | 0.403 |
| TBCD | ZMAT5 | 0.341 |

| Top 3 correlated biomarkers in sets S1 and S2 |  |  |
| --- | --- | --- |
| Biomarker (S1) | Biomarker (S2) | Correlation |
| KRT26 | ZNF277 | 0.365 |
| GUCY1A3 | ZNF277 | 0.287 |
| KCTD5 | EXOSC6 | 0.286 |

#### 3.3. Comparison with other studies

It is important to compare our results with the studies conducted in the past. There has been a study in the past for identification of salivary biomarkers for Gastric Cancer. They identified a five biomarker panel (SPINK7, SEMA4B, PPL, miR-140- 5p and miR-301a-3p) for the classification of GC vs Normal and tested them on different cohorts using microarray. The detailed comparison of the studies is given in Table 7. The highest reported AUROC in this study is 0.87 on 5 biomarkers and demographics by Li et al. Unfortunately, dataset used in this study did not provide information on demographics. However, for our study, 8 mRNA biomarkers alone are able to distinguish between the two classes with an AUROC of 0.905. In order to check if these biomarkers performed well on the dataset used in this study, we used these 5 biomarkers as features and developed ML models on them. It was observed that the highest AUROC obtained on our dataset using these biomarkers was only 0.563 for independent validation set. We also attempted to add these 5 biomarkers to our set of 8 primary biomarkers to see if it improved our performance, however, this decreased the AUROC from 0.905 to 0.830 for independent validation set. These results are given in detail in Table 8.

**Table 7:**
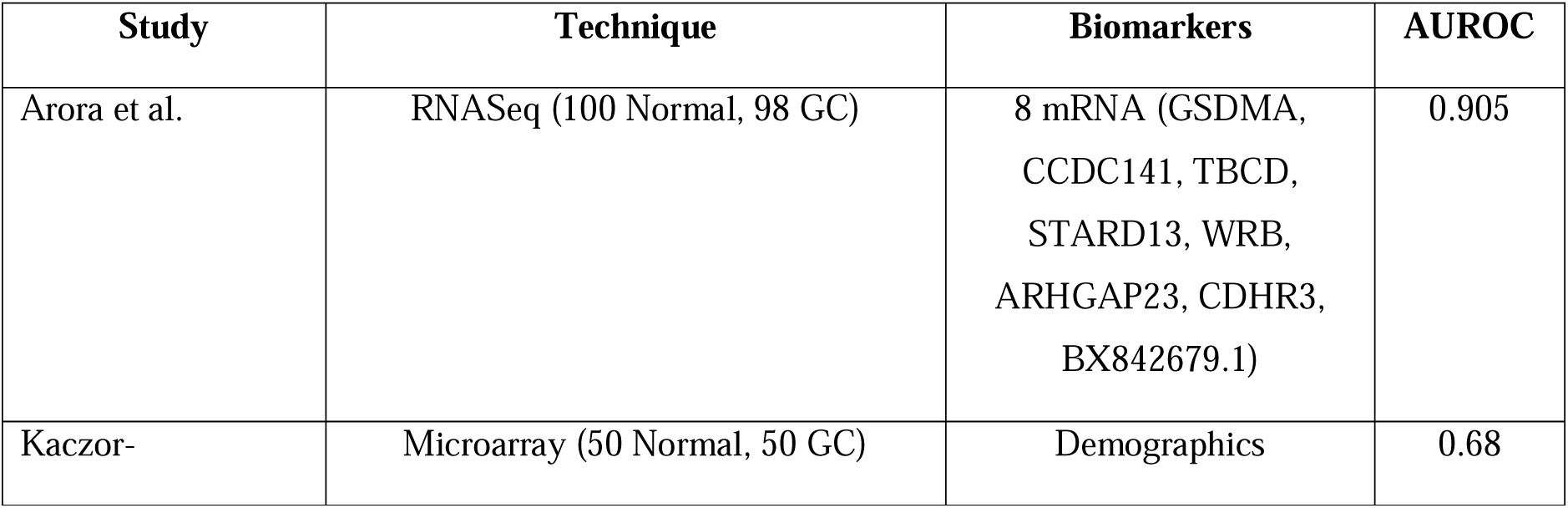
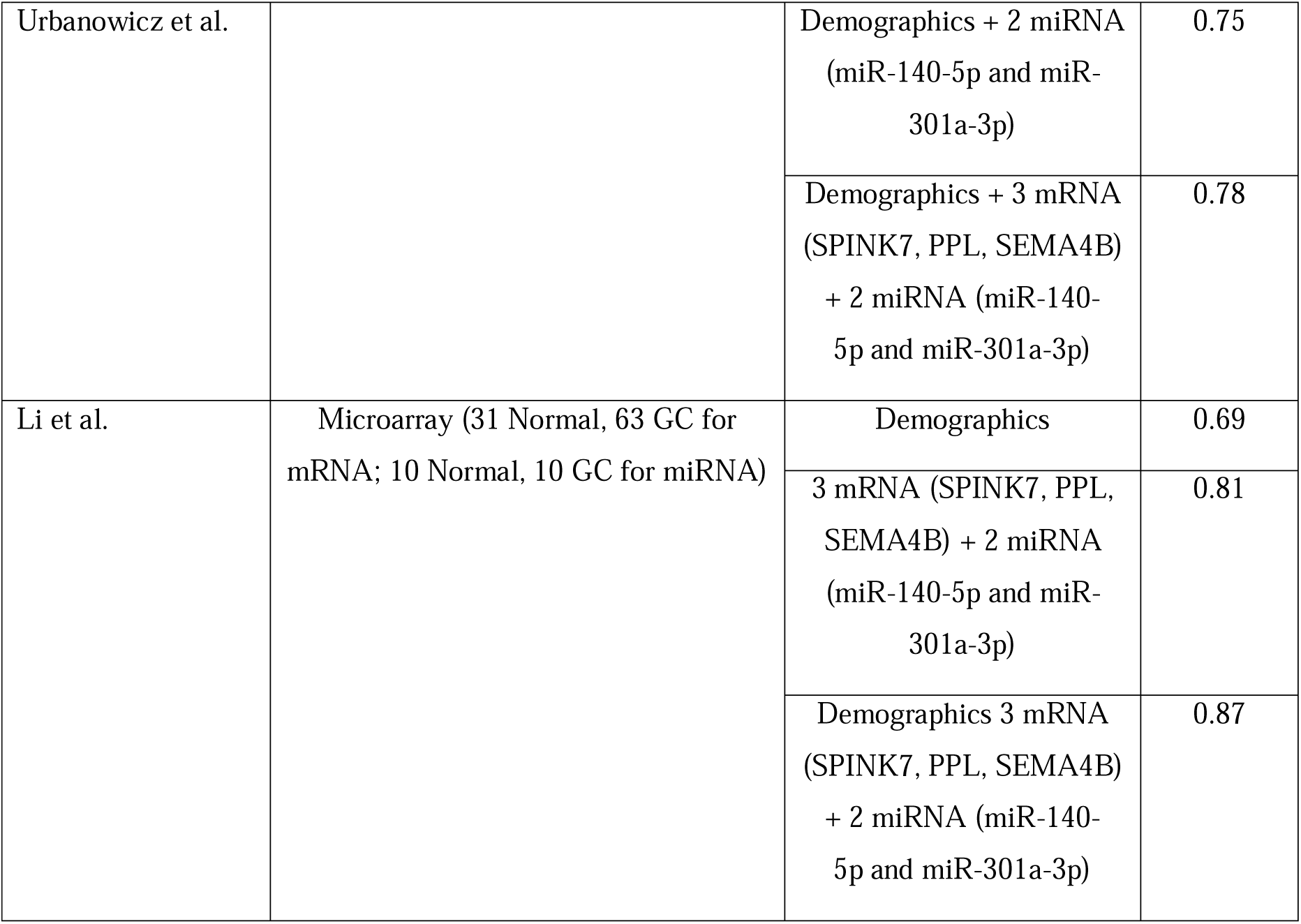
Comparative analysis of biomarker performance between our study and previous studies.

**Table 8:** Results for ML models developed on a) 5 biomarker panel discovered in Li. et al. study, b) 13 biomarkers - 8 primary biomarkers identified in this study, and 5 biomarkers identified in Li et al. study.

| Li et al. biomarkers (n=5) |  |  |  |  |  |  |  |  |  |  |  |
| --- | --- | --- | --- | --- | --- | --- | --- | --- | --- | --- | --- |
|  |  | Training Set |  |  |  |  | Validation Set |  |  |  |  |
| Model | Thr | Sens | Spec | Acc | MCC | AUC | Sens | Spec | Acc | MCC | AUC |
| ET | 0.48 | 0.49 | 0.45 | 46.84 | -0.063 | 0.455 | 0.35 | 0.55 | 45.00 | -0.102 | 0.453 |
| RF | 0.5 | 0.51 | 0.51 | 51.27 | 0.025 | 0.499 | 0.45 | 0.75 | 60.00 | 0.210 | 0.510 |
| LR | 0.48 | 0.55 | 0.54 | 53.27 | 0.032 | 0.542 | 0.60 | 0.50 | 55.00 | 0.101 | 0.563 |
| XGB | 0.48 | 0.47 | 0.50 | 48.73 | -0.026 | 0.438 | 0.35 | 0.45 | 40.00 | -0.201 | 0.464 |
| KNN | 0.01 | 0.45 | 0.51 | 48.10 | -0.039 | 0.481 | 0.50 | 0.50 | 50.00 | 0.000 | 0.500 |
| SVC | 0.49 | 0.49 | 0.51 | 50.00 | 0.000 | 0.479 | 0.75 | 0.20 | 47.50 | -0.060 | 0.467 |
| Primary biomarkers + Li et al. biomarkers (n=13) |  |  |  |  |  |  |  |  |  |  |  |
|  |  | Training Set |  |  |  |  | Validation Set |  |  |  |  |
| Model | Thr | Sens | Spec | Acc | MCC | AUC | Sens | Spec | Acc | MCC | AUC |
| **ET** | 0.51 | 0.78 | 0.78 | 77.85 | 0.557 | 0.809 | 0.65 | 0.85 | 75.00 | 0.510 | 0.808 |
| **RF** | 0.49 | 0.72 | 0.71 | 71.52 | 0.430 | 0.789 | 0.70 | 0.80 | 75.00 | 0.503 | 0.779 |
| **LR** | 0.49 | 0.74 | 0.70 | 72.15 | 0.444 | 0.808 | 0.75 | 0.70 | 72.50 | 0.451 | 0.803 |
| **XGB** | 0.49 | 0.65 | 0.66 | 65.82 | 0.316 | 0.736 | 0.65 | 0.75 | 70.00 | 0.402 | 0.742 |
| **KNN** | 0.41 | 0.81 | 0.66 | 73.42 | 0.475 | 0.788 | 0.70 | 0.75 | 72.50 | 0.451 | 0.794 |
| **SVC** | 0.5 | 0.77 | 0.76 | 76.58 | 0.532 | 0.828 | 0.70 | 0.85 | 77.50 | 0.556 | 0.830 |

## Discussions

Gastric cancer (GC) remains a significant global health burden due to its high mortality rate and late-stage diagnosis [33]. The conventional diagnostic methods such as endoscopy and histopathology are invasive, costly, and not suitable for routine population screening [34]. In this context, saliva has emerged as a highly promising biofluid for non-invasive biomarker discovery. Saliva contains a diverse repertoire of extracellular molecules, including proteins, metabolites, DNA, and RNA, that can reflect systemic physiological and pathological changes [6,35,36].

In this study, we have attempted to identify salivary extracellular RNA (exRNA) biomarkers that can differentiate between Gastric Cancer (GC) and Normal samples accurately. For this, we firstly identified statistically significant (p-value<0.05) salivary exRNA by applying Mann-whitney U test on 98 GC and 100 Normal samples containing read counts for ∼99,000 salivary exRNA. Then, we performed a threshold-based classification of GC vs Normal using the mid-point of the sum of means of both populations as a threshold. Through this, we identified that biomarker GDSMA (downregulated) was able to classify GC vs Normal with an accuracy of 64.14%, followed by SMR3B (Upregulated) with an accuracy of 62.63%, and then NPTXR (Upregulated), KCNC4 (Downregulated), TBCD (Upregulated), and KRT26 (Downregulated) with accuracies of 61.61%.

Next, to enhance the prediction of GC vs Normal samples, we applied Machine Learning (ML) models to the dataset. We divided the data into 80% training and 20% independent validation sets. Firstly, we wanted to develop a gene-based classification model, hence we selected only genes (n=340) out of the significantly different salivary exRNA. The LR model developed on these genes achieved an AUC of 0.975 on an independent validation set. On further performing feature selection methods on this set, we identified 9 genes (GSDMA, HSP90B1, SLC30A4, ATP8B3, ARHGAP23, NPTXR, WRB, SMR3B) which achieved the highest AUC of 0.893 using voting classifier.

To further improve the prediction, and identify other relevant exRNA apart from mRNA, we included the most common categories of RNA that are used as biomarkers - circRNA, miRNA, piRNA, tRNA, along with mRNA. This data consisted of 362 salivary exRNA which achieved an AUC of 0.965 and MCC of 0.851 for Logistic Regression (LR) Model. However, we wanted to identify top biomarkers that could differentiate between GC and Normal with high accuracy. Using feature selection techniques, we were able to identify a biomarkers set containing 24 biomarkers (23 genes and 1 tRNA) which had comparable performance with an AUROC of 0.937 and MCC of 0.851 on independent validation dataset. However, we wanted to further reduce the number of biomarkers making it more feasible to be used in clinical studies, which led to identification of 8 biomarkers (GSDMA, CCDC141, TBCD, STARD13, WRB, ARHGAP23, CDHR3, BX842679.1) using feature selection techniques with the highest AUROC of 0.905 and MCC of 0.770 using ensemble classifier – Stacking classifier, which surpassed the performance of the set selected using just mRNA.

Feature selection techniques aim to enhance model performance by selecting the most informative features while reducing redundancy [37]. When faced with highly correlated features, many selection methods tend to retain only one representative feature to avoid multicollinearity and redundant information. In biomarker identification, blindly removing one feature from a set of highly correlated biomarkers can be problematic because correlated features may still hold distinct biological significance. To combat this issue, we attempted to identify multiple significant set of biomarkers. We named the biomarker set having 8 biomarkers that was identified first as primary set (P) of biomarkers. To identify more relevant biomarkers, we firstly removed this set from the initial set of 362 biomarkers (A), and then performed feature selection again to identify first secondary set (S1) of biomarkers that contained 11 salivary exRNA (KRT26, GUCY1A3, KCTD5, CADPS, ATR, ALX1, TMEM98, HIST1H3A, KPTN, hsa_circ_000353, ZNF688). We then repeated this process to obtain the second secondary set of biomarkers (S2) which also contained 11 salivary exRNA (KIAA2022, PABPC3, MRPL51, CALB2, MDH1B, ZNF277, PRKD3, EXOSC6, LPPR5, ATP8B3, ZMAT5). We also calculated correlation of these identified biomarkers in each set with each other. It was observed that in sets P and S1, the highest correlated biomarkers in these sets were GSDMA with GUCY1A3 with a Pearson’s correlation of 0.529. For sets P and S2, highest correlation was amongst ARHGAP23 and LPPR5 having the value of 0.411, and for sets S1 and S2, highest correlation was amongst mRNA biomarkers KRT26 AND ZNF277 with a value of 0.365.

We compared the performance of our ML model with the existing studies, and it was observed that our set of biomarkers outperformed the existing studies, which report the highest AUC of 0.87 for 3 mRNA and 2 miRNA biomarkers with demographics. When we used these biomarkers on our dataset to classify GC vs Normal, the highest AUROC was only 0.563 on the independent validation dataset. On adding these biomarkers to our primary set of biomarkers (P), our AUROC performance reduced from 0.905 to 0.830 on the independent validation dataset.

The primary set of biomarkers identified in this study – GSDMA (downregulated), CCDC141 (downregulated), TBCD (upregulated), STARD13 (downregulated), WRB (downregulated), ARHGAP23 (upregulated), CDHR3 (upregulated), BX842679.1 (downregulated), have been previously observe to have roles in Gastric Cancer pathogenesis. GSDMA regulates apoptosis in gastric epithelial cells and been observed to be frequently silenced in GC [38,39]. CCDC141 has also been previously reported to be involved in cellular processes related to cancer progression and metastasis and also shown to be associated with GC[40]. STARD13 has been reported to function as a tumor suppressor, and STARD13 and NEU1 are reported to be direct targets of miR-125b in GC [41]. CDH5 has been previously observed to be involved in the regulation of multiple cancer targets and can act as a prognostic biomarker in GC [42]. ARGHAP23 regulates actin cytoskeleton and cell motility and is often seen upregulated in GC [43]. CDHR3 has been previously linked to cell adhesion processes and is seen dysregulated in GC [44,45]. However, for BX842679.1, there is limited information on its function in GC but it is found to be associated with Myeloid Leukaemia [46]. TBCD is involved in cellular proliferation and microtubule dynamics, but no association with GC mechanism has been reported [47].

A key limitation of this study on GC salivary exRNA biomarkers is that it is primarily computational, relying on bioinformatics-driven analysis of existing datasets. While the findings provide valuable insights into potential biomarker candidates, they should be interpreted as preliminary rather than definitive. To establish their true diagnostic and prognostic value, further wet-lab validation is required.

## Author’s contributions

AA retrieved and processed the data, implemented the algorithms, and developed the prediction models. AA, and GPSR prepared the manuscript. GPSR conceived and coordinated the project. All authors have read and approved the final manuscript.

## Acknowledgements

The authors are thankful to the Council of Scientific and Industrial Research (CSIR) for providing the fellowship. The authors are also thankful to the Department of Computational Biology, IIITD, New Delhi for its infrastructure and facilities. We thank the Department of Biotechnology (DBT) for providing an infrastructure grant to the institute (Grant BT/PR40158/BTIS/137/24/2021). We would like to acknowledge that figures were created using BioRender, and English was corrected using Grammarly.

